# Evidence that stress-induced changes in surface temperature serve a thermoregulatory function

**DOI:** 10.1101/788182

**Authors:** Joshua K. Robertson, Gabriela Mastromonaco, Gary Burness

## Abstract

Changes in body temperature following exposure to stressors have been documented for nearly two millennia, however, the functional value of this phenomenon is poorly understood. We tested two competing hypotheses to explain stress-induced changes in temperature, with respect to surface tissues. Under the first hypothesis, changes in surface temperature are a consequence of vasoconstriction that occurs to attenuate blood-loss in the event of injury and serves no functional purpose *per se;* defined as the Haemoprotective Hypothesis. Under the second hypothesis, changes in surface temperature reduce thermoregulatory burdens experienced during activation of a stress response, and thus hold a direct functional value; here, the Thermoprotective Hypothesis. To understand whether stress-induced changes in surface temperature have functional consequences, we tested predictions of the Haemoprotective and Thermoprotective hypotheses by exposing Black-capped Chickadees (n = 20) to rotating stressors across an ecologically relevant ambient temperature gradient, while non-invasively monitoring surface temperature (eye region temperature) using infrared thermography. Our results show that individuals exposed to rotating stressors reduce surface temperature and dry heat loss at low ambient temperature and increase surface temperature and dry heat loss at high ambient temperature, when compared to controls. These results support the Thermoprotective Hypothesis and suggest that changes in surface temperature following stress exposure have functional consequences and are consistent with an adaptation. Such findings emphasize the importance of the thermal environment in shaping physiological responses to stressors in vertebrates, and in doing so, raise questions about their suitability within the context of a changing climate.

**Summary Statement:** We provide empirical evidence for a functional value to stress-induced changes in surface temperature that is consistent with an adaptation, using a temperate endotherm (Black-capped Chickadee) as a model species.

## Introduction

Changes in body temperature following perception of a stressor have been known for nearly two millennia. In the 2nd centurey CE, for example, Galen described the presence of an “ephemeral fever” in humans that was thought to be evoked by an abundance of humour (Yeo, 2005); a concept later expanded upon by Ibn Sina (11th century CE) who described “fevers” driven by emotions, including grief, anger, and dread (Avicenna, 2005; Parviz et al, 2013). To date, changes in body temperature following exposure to stressors have been reported across numerous non-hominid species (i.e. lizards: Cabanac and Gosselin, 1993; turtles: Cabanac and Bernieri, 2000; birds: Greenacre and Lusby, 2004, and fish: Rey et al, 2015) and garnered significant research attention. Despite such long-standing recognition in philisophical and scientific literature, and near vertebrate-wide conservation (but see Cabanac and Laberge, 1998; Jones et al, 2019), the functional significance of stress-induced changes in body temperature remain elusive. Indeed, while many studies have done well to uncover proximate mechanisms underpinning this phenomenon (i.e. mediation by peripheral vasoconstriction, pyrogenic cytokines, prostaglandins, or glucocorticoids; Yokoi, 1966; reviewed in Oka et al, 2001; and Jerem et al, 2018, respectively), remarkably few have empirically interrogated a functional role of changing one’s body temperature, *per se*, in response to a stress exposure (Cabanac and Gosselin, 1993).

Thermal responses to stress exposure can be broadly categorized according to their position of occurrence within an organism (i.e. at core tissues, or surface tissues). Of such positions, core temperature responses have, perhaps, received the greatest theoretical attention with respect to functional significance (reviewed in Oka et al, 2001; Oka, 2018). For example, some immunological studies have posited that changes in core body temperature following stress exposure represent a true “fever” (Singer et al, 1986; Sanches et al, 2002), and endow individuals with a defensive advantage if they experience injury and pathogen exposure during stress exposure (discussed in Oka et al, 2001). Others, however, have contested this hypothesis by failing to substantiate immune-mediation of core temperature responses to stress (Long et al, 1990b, Soszynski et al, 1998; Hiramoto et al, 2009; Vinkers et al, 2009). At the level of surface tissues, however, a functional role of stress-induced changes in temperature, *per se*, appears yet to be raised (though breifly discussed in Herborn et al, 2018). Indeed, dominant theory explaining stress-induced changes in surface temperature posits that this phenomenon is merely a consequence of haemetic redistribution (Jerem et al, 2015; Jerem et al, 2018, Nord and Folkow, 2019) and holds no direct functional role; rather, it is haemetic redistribution, but not thermal modulation, that carries functional significance by attenuating blood loss in the event of injury (as long-shown following hemorrhage in McGuigan and Atkinson, 1921; Freeman, 1932; Darlington et al, 1986).

Despite an absence of functionally guided research, studies describing stress-induced changes in surface temperature do allude to a functional value of this phenomenon. Nord and Folkow (2019), for example, reported that while skin temperature of Svalbard Rock Ptarmigans (*Lagopus muta hyperborea*) typically falls after handling, the magnitude of the skin temperature response varies according to ambient temperature (*Ta*); specifically, Ptarmigans handled at low temperature (-20°C) display a larger change in skin temperature than those handled at warmer temperatures (0 °C). An effect of *Ta* on stress-induced changes in skin temperature suggests that the function of this phenomenon extends beyond haemetic redistribution and may provide thermoregulatory advantages, where heat conservation at low *Ta* (sub-thermoneutral) is enhanced during perception of environmental challenges. Supporting a thermoregulatory function to stress-induced changes in surface temperature, Herborn et al (2018) described an increase in skin temperature of Domestic Chickens (*Gallus gallus*) after exposure to chronic stress treatments at constant, thermoneutral temperatures, with respect to controls. Given exposure to stressors is thought to elevate metabolically generated heat (i.e. as a consequence of tachycardia, tachypnea, and avoidance behaviour; Cabanac and Aizawa, 2000; Cabanac and Guillemette, 2001; Greenacre and Lusby, 2004; Long et al, 1990a), elevation of skin temperature under chronically challenging environments may facilitate dissipation of excess metabolically-generated heat, thereby reducing thermal load associated with activation of a stress response. Neither Nord and Folkow (2019), nor Herborn et al (2018), however, tested patterns of heat conservation or dissipation following stress exposure, rendering such thermoregulatory consequences of surface temperature responses speculative.

In this study, we propose that stress-induced changes in surface temperature, *per se*, serve a functional role that may be understood when contextualized according to energetic and thermal load. Specifically, we argue that changes in surface temperature following stress exposure reduce energetic costs that are incurred during activation of a stress response, by promoting heat conservation at low temperatures (conservatively, below thermoneutrality), and heat dissipation at high temperature (conservatively, above thermoneutrality). This hypothesis (henceforth defined as the “Thermoprotective Hypothesis”) contrasts the dominant hypothesis stating that surface temperature responses are a functionally neutral corollary of haemetic redistribution (discussed above, and in Jerem et al, 2015; henceforth defined as the “Haemoprotective Hypothesis”).

To test whether stress-induced changes in surface temperature themselves provide a functional value, we generated predications for both the Thermoprotective Hypothesis and Haemoprotective Hypothesis, with reference to a temperature endotherm, the Black-capped Chickadee (*Poecile atricapilus;* Linnaeus, 1766). According to the Thermoprotective Hypothesis, we predicted that surface temperature and dry heat-loss (here, heat lost by radiation and conduction, but not evaporative cooling) of Chickadees would fall under stress exposure when *Ta* is low (below thermoneutrality), and rise when *Ta* is high (above thermoneutrality), thereby reducing energetic and thermal load during activation of a stress response. Alternatively, under the Haemoprotective Hypothesis, we predicted that surface temperature and dry heat-loss of Chickadees would also fall under stress exposure, however, only until *Ta* and surface temperature are approximately equal (i.e. when acquisition of heat at surface tissues matches loss of heat from local ischemia). Because Black-capped Chickadees are sexually size-dimorphic (Foote et al, 2010), and body size is thought to influence both thermal conductance and heat dissipation capacity in endotherms (e.g. Porter and Kearney, 2009; Alonso et al, 2016), we further predicted that the thermal responses to stress exposure would differ between females and males under the Thermoprotective Hypothesis, but not the Haemoprotective Hypothesis. Specifically, if stress-induced changes in surface temperature follow the Thermoprotective Hypothesis, we predicted that male Chickadees, being larger than female Chickadees and having lower heat dissipation capacity, would exhibit a more robust change in surface temperature and dry heat loss across a *Ta* gradient following stress exposure than females; that is, the slope of the relationship between *Ta* and surface temperature or dry heat-loss would be steeper in males than in females, following stress exposure. If stress-induced changes in surface temperature follow the Haemoprotective Hypothesis, however, we predicted that surface temperature and dry heat loss of female and male Chickadees would respond equivalently to stress exposure across a *Ta* gradient.

To our knowledge, this is the first study to empirically test a functional role of stress-induced changes in surface temperature *per se.* It is also the first study to investigate stress-induced changes in surface temperature across an ecologically relevant *Ta* range that extends both below and above the thermoneutral zone of the study organism. In light of a changing global climate, elucidating a thermoregulatory function to the vertebrate stress response may lend insight into the adaptive capacity of organisms that face both physiological and thermal stress from anthropogenic, environmental change.

## Methods

All methods used for animal capture, handling, and experimentation were approved by the Trent University Animal Care Committee (AUP # 24614) and Environment and Climate Change Canada (permit# 10756E).

### Black-capped Chickadee Capture, Sampling and Transport

During the months of March and April in 2018, we captured 20 free-living Black-capped Chickadees within a 100 km^2^ area of south-central Ontario (Canada) for captive experimentation. Because conspecific songbirds from urban and rural populations have been shown to exhibit differences in the magnitude of the stress response (e.g. Abolins-Abols et al, 2016), 10 individuals (n_females_ = 5; n_males_ = 5) were captured from across known urban populations (Cambridge, Ontario; 43.3789° N, 80.3525° W, Guelph, Ontario; 43.3300° N, 80.1500° W, Brantford, Ontario; 43.1345° N, 80.3439° W) and 10 (n_females_ = 5; n_males_ = 5) were captured across three known rural populations (Erin, Ontario; 43.7617° N, 80.1529° W, Corwhin, Ontario; 43.5090° N, 80.0899 ° W, Ruthven Park National Historic Site, Ontario; 42.9797° N, 79.8745° W). All Chickadees were trapped using remotely operated potter-traps (90.0 cm × 70.0 cm × 70.0 cm; 1 × w × h), baited with sunflower seeds and suet on the day of capture. To further attract individuals to trapping locations, Black-capped Chickadee breeding and mobbing songs were broadcasted alternately from a remote call-box (FoxPro™ Patriot; Lewisville, Pennsylvania, USA) at approximately 80 decibels, then ceased when at least one individual had approached within a four-meter radius of a baited trap.

Once captured, individuals were immediately blood sampled (50μL) by brachial venipuncture and capillary tube collection (< 5 minutes post-capture), then assigned a unique combination of one government issued, stainless steel leg band, and two coloured, darvic leg band for identification. Individuals were then weighed (nearest 0.1 g using a digital platform scale), measured (tarsus (mm) and flattened wing-chord (mm) to the nearest 0.1 mm, using analogue calipers) and secured in covered carrier cages (30.0 cm × 30.0 cm × 15.0 cm; 1 × w × h) for transport to our long-term holding facilities at the Ruthven Park National Historic Site, Cayuga, Ontario, Canada (maximum of 90 km; 2 hours travel by vehicle). Finally, erythrocytes were isolated from whole blood samples on site by centrifugation (12,000 rotations per minute), then lysed in 500μL Queen’s Lysis Buffer for long-term preservation of genetic material (Seutin et al, 1991). Plasma was isolated and stored for another study (not described here). All lysed blood samples were held on ice until transfer to permanent holding at 4°C.

### Experimental Enclosures and Maintenance

Once at our long-term holding facility, all Chickadees were randomly assigned to one of four identical and visually isolated outdoor flight enclosures (n = 5 per flight enclosure; 90 cm × 60 cm × 200 cm; 1 × w × h). Each flight enclosure was supplied with an insulated roosting box (60 cm × 20 × 20 cm; 1 × w × h) mounted at 1.2 m in height, one roosting tree (White Cedar, *Thuja occidentalis;* 1.0 m), and two perching branches (80 cm in length), mounted at approximately 1.5 and 1.8 m in height. Chickadees were provided water, and a mixture of meal worms (*Tenebrio molitor*), house crickets (*Acheta domesticus*), shelled peanuts, apple pieces, boiled egg, sunflower seeds, safflower seeds, and Mazuri™ (St. Louis, Missouri, USA) Small Bird Maintenance diet *ad libitum.* Both food and water were distributed twice to three times daily, with food being exclusively dispensed on a 400 cm^2^ platform, raised to 1.2 m in height. To minimize disturbance during feeding, all food and water were distributed through opaque, hinged doors (15 cm × 15 cm), such that Chickadees were blind to experimenter presence. All individuals were acclimated for a minimum of two weeks prior to experimentation.

### Experimental Stress Induction

From April until late June of 2018, we tested a functional role of stress-induced changes in surface temperature by using a repeated sampling design to maximize statistical power. Here, each individual was exposed to one control (untouched; n = 30 days) and one stress exposure treatment (n = 30 days), separated by a rest period of 2 days (total experimental duration = 62 days). All Chickadees were maintained in outdoor flight enclosures for the duration of the study (see “Experimental Enclosures and Maintenance”) and were therefore exposed to seasonal changes in temperature and day length that could alter patterns of stress responsiveness (Wingfield et al, 1992; Astheimer et al, 1995). To account for these seasonal changes, individuals within two flight enclosures (n = 10; Group A) were exposed to a control treatment followed by a stress exposure treatment, while the remaining individuals (n = 10 within two flight pens; Group B) were exposed to a reversed treatment order, such that stress exposure treatments in Group B co-occurred with control treatments in Group A, and control treatments in Group B co-occurred with stress exposure treatments in Group A.

Stress exposure treatments followed a protocol of rotational stressors similar to Rich and Romero (2005) and Cyr and Romero (2007), however, no auditory stressors were used to ensure that the application of a stressor to individuals within one flight enclosure did not elicit a stress response in individuals held within remaining and nearby flight enclosures. Here, experimental individuals were administered five, randomly selected passive stressors each day, with each stressor persisting for 20 minutes, and being separated from subsequent stressors by 1 hour. Daily randomization of stressors was used to circumvent habituation to each individual stressor throughout the course of experimentation. Stressors included; capture and restraint, presence of a mock predator (adult Cooper’s Hawk, *Accipiter cooperii*), presence of a novel object placed in the centre of holding enclosures (garden gnome), presence of a human within holding enclosures, absence of light (by enwrapping enclosures with opaque fabric), and presence of a mounted conspecific placed in the centre of a feeding platform, mimicking an unfamiliar and dominant individual. Hormonal responses to stress exposure treatments were not measured since blood collection is likely to interfere with peripheral thermal profiles, however, behavioural responses to each stressor were visually and statistically confirmed (alarm calling, panting, and elicitation of avoidance behaviour during stress exposure; JKR personal obs; reduction in feeding; Supplemental Results and SFig. 1). Individuals within control treatments were maintained according to acclimation conditions and were not subject to handling or disturbance by experimenters.

**Figure 1.**
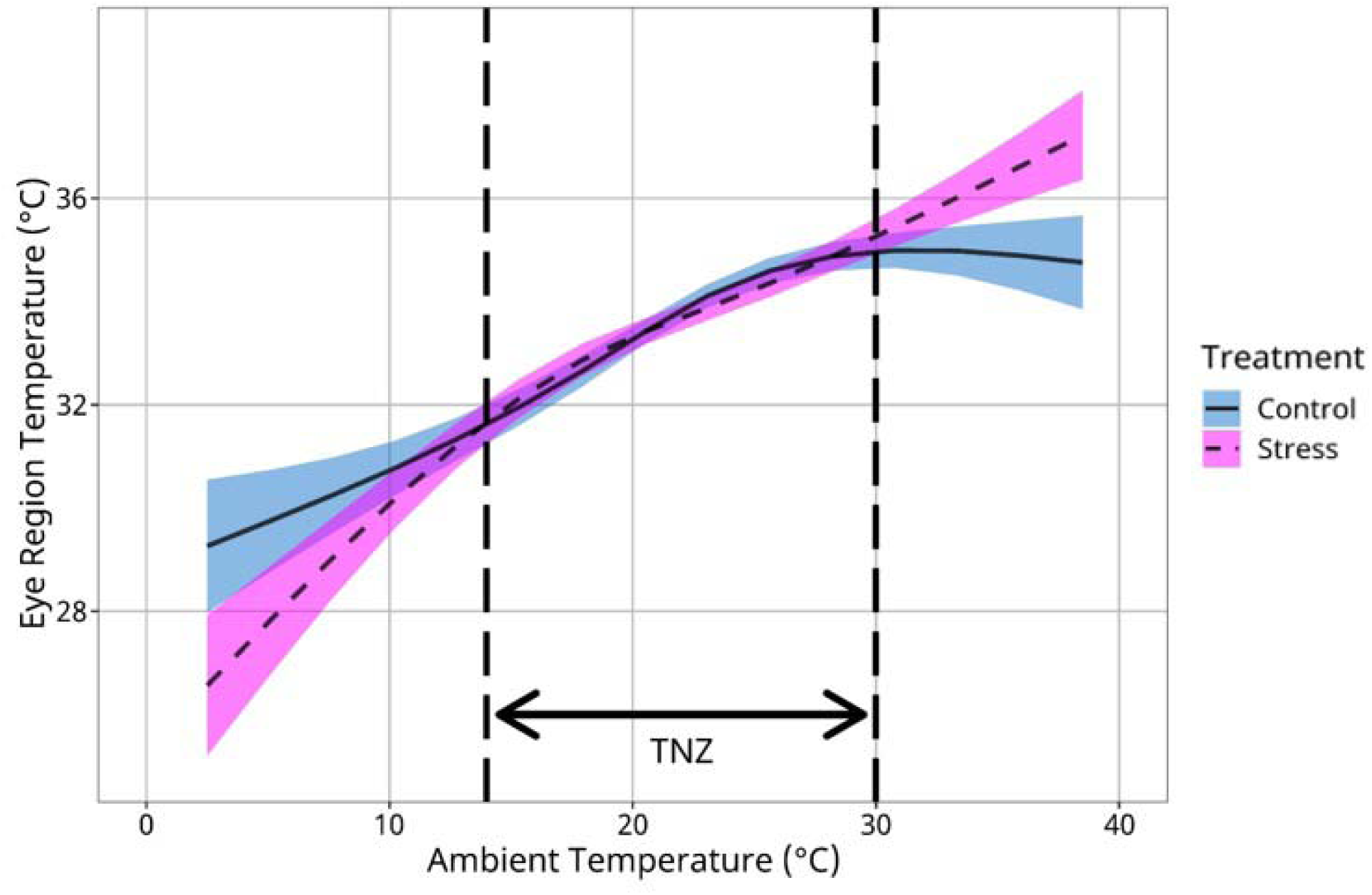
Eye temperature (°C) of Black-capped Chickadees exposed to repeated stressor and control treatments (n = 20 individuals, n = 30 days per treatment), according to *Ta* (°C). Eye temperature values are derived from thermographic images (n = 6431) captured during feeding. Trend-lines represent the estimated marginal means of eye temperature according to *Ta*, as determined from a generalized additive mixed effects model (“GAMM”). Blue shadows represent 95% confidence intervals around means for control treatments, and magenta shadows represent 95% confidence intervals around means for stressed treatments. Vertical dashed lines represent limits to the thermoneutral zone (or “TNZ”).

At the onset and completion of each treatment (control and stress exposure), all individuals were re-weighed to monitor changes in body condition across treatments (see Supplemental Methods and Results), and upon completion of the experiment, individuals were released to their site of capture.

### Environmental Data Collection

We were interested in testing the effects of repeated stress-induction on surface temperature profiles, within the context of naturally cycling temperatures. We therefore sampled *Ta* (°C) at the location of experimentation using a ThermoChron iButtonTM (Maxim Integrated, DS1922L-F5; San Jose, California, USA) placed in the shade, for the duration of experimentation. To capture rapid and subtle changes in *Ta* across time, *Ta* readings were sampled at a frequency of 20 samples/hour, and at a resolution of 0.5°C. Because relative humidity can influence the transmission of infrared radiation, and therefore estimates of object temperature by thermography (reviewed in Tattersall, 2016), we collected local relative humidity in addition to *Ta* (one sample/hour) from Environment Canada climate repositories (https://climate.weather.gc.ca/; station - Hamilton A; 22 km from the experimental holding location).

### Thermographic Filming

Surface temperature responses to stress exposure and control treatments were measured by capturing maximum eye region temperature of Chickadees during feeding, using time-lapse infrared thermography. Temperature of the eye region was assessed because it contains exposed integument that may be readily imaged in birds while stationary (e.g. during feeding) and is capable of heat-exchange unfettered by insulatory keratinous tissues (i.e. feathers or leg-scale; discussed in Jerem et al, 2018). Furthermore, in Domestic Chickens, exposure to hyperthemic conditions has been shown to increase blood-flow in capillary beds of the head (Wolfensen et al, 1981), suggesting that cephalic vasculature, including the vessels located near the eye region (i.e. the opthalmotemporal, ethmoid, and facial veins), may serve as a location for heat exchange (discussed in Midtgård, 1983). Here, we chose to use the maximum temperature of the eye region as a metric of eye region temperature, rather than mean temperature, because it is thought to be less susceptible to measurement error (Jerem et al, 2015; Jerem et al, 2018), and is less likely to fluctuate according to the angle at which an individual was imaged.

Thermographic images were captured using a remotely operated thermographic camera (VueProR™, FLIR, Wilsonville, Oregon, USA; 13 mm lens, 336 × 256 resolution; image frequency = 1 Hz) that was rotated among flight enclosures each day, according to cardinal direction. Specifically, beginning at 08:00 each day, time-lapse thermographic imaging was conducted for approximately 1 hour at one flight enclosure, after which the thermographic camera was transferred cardinally clockwise to a second flight enclosure, and imaging was repeated for 1 hour. This rotational process was repeated until 16:00 each day, to ensure that each flight enclosure was subjected to at least one hour of thermographic filming per day. To control for possible effects of circadian rhythms on surface temperature profiles, the first flight enclosure to be filmed each day was also rotated cardinally clockwise direction. Because individuals within a flight enclosure could not be identified by thermography (according to colour-band combinations), a remotely operated digital camera was rotated alongside the thermographic camera, allowing for *post hoc* individual identification.

To ensure that Chickadees were blind to the presence of both the thermographic and digital camera during experimentation, flight enclosures were equipped with water-tight camera boxes that were mounted to an exterior wall adjacent to the feeding platforms, and were perforated with two 30 mm diameter holes through which thermographic imaging and digital filming were conducted (distance of 0.5 m from feeding platforms). Camera boxes were solely accessed from the exterior of flight enclosures, where an experimenter could not be seen by Chickadees within the flight enclosure. When thermographic imaging was not being conducted, both 30 mm holes were covered.

### Data Extraction from Thermographic Images

Throughout experimentation, 1,035,512 thermographic images were captured across 60 days. Raw radiance values (kW/m^2^) per pixel were extracted from all thermographic images in R (version 3.6.1), then converted to temperature values (°C) using Planck’s law, and following equations described by Minkina and Dudzik (2009) and Tattersall (2016). Here, temperature values were calibrated according to *Ta* and relative humidity (determined in “Environmental Data Collection”), and calibration and atmospheric constants for our thermographic camera were identified using Exiftool (Harvey, 2018). Emissivity of the eye region was assumed to be 0.95, according to estimates for avian integument by Best and Fowler (1981).

Following conversion of radiance to temperature in thermographic images, we used FIJI (Schindelin et al, 2010) to determine maximum eye region temperature (°C) for Chickadees that were present in thermographic images by drawing a region of interest (ROI) around the eye region (approximately 1.0 cm in diameter) and extracting the maximum temperature from within the ROI. Because motion of an object can lead to underestimation of its surface temperature in thermographic imaging (discussed in Jerem et al, 2018), only images where the feeding individual was stationary (i.e. not in flight, landing from flight, or departing) were included in our analyses. Furthermore, maximum eye region temperature was only selected from images where the identity of the individual imaged could be determined from parallel digital video (n = 6431; n_Stress_ = 3397, n_Control_ = 3034).

### Heat Transfer Calculations

The Thermoprotective Hypothesis predicts that stress-induced changes in surface temperature alleviate thermal burdens incurred during a stress response, with stressed individuals conserving more heat at low *Ta* and dissipating more heat at high *Ta* than control individuals. To test this prediction, we sought to quantify and compare total heat transfer (q_tot_) from the eye region of captive Chickadee, in stress-induced and control treatments. Here, heat transfer was calculated from thermographic images using methods described by Ward et al (1999), McCafferty et al (2011), and Nord and Nilsson (2019).

Because individuals were sheltered from wind in flight enclosures, and were unlikely to transfer heat by direct contact between the eye region and a medium other than air, q_Total_ was assumed to be the sum of radiative heat transfer (q_Rad_), and free conductive heat transfer (qConv). Here, the values for kinematic viscosity of air (m^2^/S; at an assumed atmospheric pressure of 101.325 kPa), thermal conductivity of air (W/m °C^-1^), and thermal expansion coefficient of air (1/K) used in the calculation of qRad and q_Conv_ were calculated according to *Ta* at the time of image capture. Eye region surfaces were treated as planar structures, similar to the ventral surfaces of Blue Tits (*Parus caeruleus*) described in Nord and Nilsson (2019), and the surface area of the imaged eye region was estimated to be an oval of 1.0 cm vertical diameter and 1.1 cm horizontal diameter. Final q_Total_ values were multiplied by two to represent total heat transfer across both eye regions of an individual.

### DNA Extraction and Genetic Sexing

Sex of Black-capped Chickadees was determined genetically according to Griffiths et al (1996) and Fridolfsson and Ellegren (1999). Briefly, whole DNA was isolated from lysed erythrocyte samples by phenol:chloroform:isoamyl alcohol (25:24:1; Fisher Scientific, Waltham, Massachusetts, USA) extraction and precipitation in 100% 2-propanol, then stored at -20°C. Following DNA isolation, sex of individuals was determined by PCR amplification of chromohelicase-DNA-binding-protein intron 16 (Griffiths et al, 1996; Fridolfsson and Ellegren, 1999), and size separation of amplicons on 3% agarose gels.

### Statistical Analyses

All statistical analyses were conducted in R version 3.6.1 (R Core Team, 2019) with *a* set to 0.05. All plots were produced using the r package “ggplot2” (Wickham, 2016).

### Effect of Stress on Surface Temperature across Ambient Temperature

To test whether *Ta* influenced how eye region temperature responded to stress exposure treatments, we used a generalized additive mixed effects model (GAMM) in the package “mgcv” (Wood, 2011), fitted with restricted extent maximum likelihood (or “REML”). Here, we chose to employ an additive model rather than a linear model to ensure that non-linear changes in surface temperature across *Ta*, that are expected in species capable of thermoregulation, were captured. In this model, maximum eye temperature (°C) was used as the response variable, and treatment (binomial; control, stress-induced) was included as a parametric predictor to test for differences in the intercept of maximum eye temperature between groups. The effect of *Ta* on eye temperature was tested by including a cubic regression spline for *Ta*, however, the number of knots used in spline construction was limited to four to both capture a curvilinear relationship and avoid model over-fit. Some Parid species have been shown to exhibit diel fluctuation in body temperature that cannot be explained by *Ta* alone (i.e. Willow Tits, *Poecile montanus* (Reinertsen and Haftorn, 1984), and Mountain Chickadee, *Poecile gambeli* (Cooper and Gessaman, 2005)). To control for variance explained by diel rhythms, time of day (seconds; midnight = 0 seconds) was also included as a cubic regression spline, with the number of knots again restricted to four. To test the influence of experimental treatment on thermal responses to *Ta* and time of day, we included interaction terms between treatment and each regression spline (*Ta* and time of day). Variance explained by individual identification, flight enclosure, and date were estimated by inclusion of random intercepts, and differences in exposure to solar radiation per flight enclosure were estimated by constructing random slopes per enclosure, by time of day. Finally, capture location (each of six locations) was included as a random intercept to account for possible physiological effects at the population level, and the possibility of relatedness among individuals from the same point of capture. Autocorrelation between adjacent time-points was corrected for by using a first order autoregressive correlation structure in “itsadug” (van Rij et al, 2017), and an estimated *φ* of 0.67. Our final predictive model was therefore as follows:

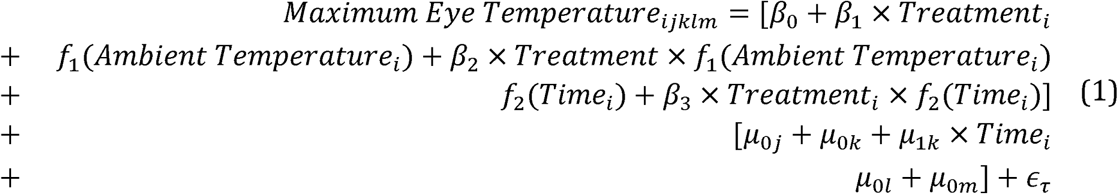

where *i* represents observation number, *j* represents individual identity, *k* represents the flight enclosure, *l* represents the date, *m* represents the capture locale, and *ε_τ_* represents a normally distributed error term.

Differences in thermal responses to stress between sexes are predicted by the Thermoprotective Hypothesis, but not Haemoprotective Hypothesis. To test whether sex influenced how surface temperature responded to stress exposure, we used a similar analytical approach to Petit and Vézina (Petit and Vézina, 2014) and first tested whether there was evidence to suggest that surface temperature responses to treatment across *Ta* differed among individuals. We did so by repeating the previous statistical model (model 1) and including both a random linear slope and a random intercept for the interaction term between *Ta* and treatment (defined as a “reaction term”), per individual Chickadee as follows;

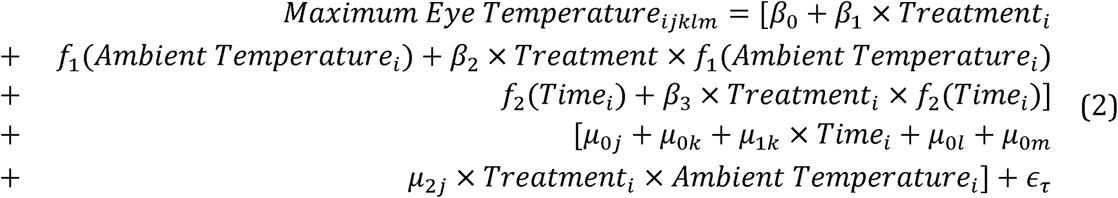

where and all terms remain as defined previously. Log-likelihood values were then calculated for our base model (model 1) and individually adjusted model (model 2), then compared using a chi-squared difference test. Our individual adjusted model (model 2) yielded a significantly higher log-likelihood, suggesting that individuals significantly differed in their surface temperature responses to stress across *Ta* (Log *𝓛*_initial_ = - 1.009×104, Log *𝓛*_adjusted_ = -9.985×103; *χ^2^=* 208.443, *df=* 28, *p <* 0.0001). We then tested whether sex could explain this individual variability by extracting the slope of the *Ta* by treatment interaction term (here, *μ*_2_) per individual and regressing them against sex as a factorial predictor in a linear model (LM; base R). Therefore, an individual’s surface temperature response to *Ta* and treatment was modeled as:

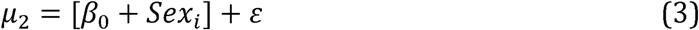

where *μ*_2_ is derived from model 2, *i* represents observation number, and *ε* represents a normally distributed error term.

### Influence of Stress Exposure on Heat Transfer across Ambient Temperature

We were interested in testing whether changes in surface temperature following stress exposure serve a thermoregulatory function. We therefore asked whether individuals exposed to repeated stressors conserved more heat below thermoneutrality, and dissipated more heat above thermoneutrality, than those exposed to control treatments. Because measurements of dry heat transfer are likely to be strongly correlated with *Ta* in endotherms (e.g. Simmons et al, 1997), we first controlled for the global effects of *Ta* on q_Tot_ by calculating how different an individual’s q_Tot_ was from that expected at a given *Ta.* This was accomplished by quantifying residual distance between observed q_Tot_ values (here, mean q_Tot_ per hour and per day, for each individual; n_Obs_ = 1006 across n_days_ = 60 and n_Birds_ = 20), and those predicted by *Ta* (mean per respective hour) using a GAMM with q_Tot_ as the response variable and mean *Ta* per hour as a cubic regression spline (restricted to four knots) in mgcv (Wood, 2011) (Mean *Ta: F=* 1.456×10^4^, *edf =* 3.000, *p <* 0.0001).

According to our Thermoprotective Hypothesis, q_Tot_ (heat lost to the environment) from an individual under stress should decrease when *Ta* is below thermoneutrality and increase above thermoneutrality. We therefore divided *Ta* measurements into temperature zones, according to their position with respect to thermoneutrality; specifically, below thermoneutrality (“low”), thermoneutral (“mid”), or above thermoneutrality (“high”). Because our experiment spanned across late winter and early summer (early April to late June; *Ta* range: 2.5°C - 38.5°C), limits of thermoneutrality in Black-capped Chickadees were generously set to 14°*C*, and 30°C, according to Grossman and West (1977) and Rising and Hudson (1974). The effect of stress-exposure on q_Tot_ across *Ta* zones was then tested using a linear mixed effect model (LMM) in the R package “lme4” (Bates et al, 2014), with residual q_Tot_ as the response variable. Treatment, temperature zone (factorial; “low”, “mid”, or “high”; total observations: n_low_ = 110, n_mid_ = 710, n_high_ = 185), and an interaction between treatment and temperature zone were included in our model as fixed effects. To test whether sex influenced the direction and magnitude of q_Tot_ across *Ta* and stress treatments (as predicted by the Thermoprotective Hypothesis), sex and a three-way interaction between sex, temperature zone, and treatment were initially included as fixed predictors in our model. Neither sex nor the three-way interaction between sex, temperature zone, and treatment were significant correlated with q_Tot_ (Sex: *p =* 0.068, Interaction between Treatment, Sex, and Temperature Zone: *p =* 0.146), and were subsequently removed to test the effects of temperature zone and treatment on q_Tot_ alone, with increased statistical power (Aiken and West, 1991). Finally, individual identity, flight enclosure, and date were included as random intercepts to account for residual variance explained by each.

## Results

### Effect of stress exposure on surface temperature and heat transfer are temperature-dependent

Eye region temperature significantly increased with *Ta* (*p <* 0.0001; Tab. 1) and this relationship was significantly influenced by treatment (*Ta*:Treatment: p = 0.0195; Tab. 1; Fig. 1), as predicted by both the Thermoprotective and Haemoprotective hypotheses. The directionality of surface temperature responses to stress, however, supported predictions of the Thermoprotective Hypothesis alone, with eye region temperature decreasing at low *Ta* and increasing at high *Ta* with respect to control treatments (Fig. 1). Interestingly, treatment alone (regardless of *Ta*) did not have a significant effect on eye region temperature (p = 0.995; Tab. 1).

**Table 1.** Results of a GAMM testing the effect of stress exposure on eye region temperature in Chickadees. Effect of *Ta* (°C), time of day (seconds), date (Julian), individual identity, and solar radiation (random slope of UT1 seconds by flight enclosure identity) are included. Estimates (*β*) are reported for parametric predictors, and estimated degrees of freedom (edf) is reported for smooth factors. Standard error values (s.e.m.) are in reference to estimates of *β* (mean *β* for smooth terms) and have been corrected for smoothness (Ʌ) in smooth terms. Asterisk (*) indicates significance at an alpha of 0.05; n_Obs_ = 6431; n_Birds_ ^=^ 20; n_Days_ = 60.

| Parametric Predictors |  |  |  |  |
| --- | --- | --- | --- | --- |
| Coefficient | Estimate ($\beta$) | s.e.m. | *t*-value | *p*-value |
| Intercept | 32.569 | 1.810 | 17.983 | <0.0001* |
| Treatment | 3.393x10<sup>-4</sup> | 0.005 | 0.007 | 0.996 |
| Smooth Predictors |  |  |  |  |
| Coefficient | *Edf* | SE | *F*-value | *p*-value |
| $Ta$ | 2.751 | 0.832 | 1.471x10<sup>4</sup> | <0.0001* |
| $Ta$ : Treatment | 2.868 | 0.797 | 7.323x10<sup>2</sup> | 0.019* |
| Time of Day | 3.451x10<sup>-5</sup> | 0.003 | <0.000 | 0.840 |
| Time : Treatment | 1.460 | 0.390 | 6.831x10<sup>1</sup> | 0.026* |
| Random Effects |  |  |  |  |
| Coefficient | SE |  |  |  |
| Individual ID | 0.161 |  |  |  |
| Flight Enclosure | 1.816 |  |  |  |
| Date | 0.425 |  |  |  |
| Time of Day \| Flight Enclosure | 1.548x10<sup>-5</sup> |  |  |  |
| Capture Location | 0.089 |  |  |  |

Although we did not detect circadian changes in eye region temperature within our observed time period (Time of day; *p =* 0.840; Tab. 1), treatment significantly influenced the relationship between time of day and eye region temperature in our birds (p = 0.026; Tab. 1). Both date of image capture and flight enclosure explained considerable variance in eye region temperature (s.e.m._Date_ = 0.425; s.e.m._Enclosure_ = 1.819; Tab. 1), probably owing to differing degrees of exposure to solar radiation per day and enclosure.

Mean q_Tot_ from the eye region was 25.275 mW (± SD = 11.903), and residual qTot from the eye region differed significantly between temperature categories (Tab. 2; Fig. 2), as predicted by the Thermoprotective Hypothesis. Specifically, individuals experiencing stress exposure conserved significantly more heat from the eye region than controls, when held below thermoneutral temperatures (p = 0.010; Tab. 2, Fig. 1), and lost significantly more heat from the eye region than control birds, when held above thermoneutral temperatures (p = 0.003; Tab. 2; Fig. 2). At thermoneutral environments, however, stress-exposed and control individuals did not differ with respect to heat exchange at the eye region (*p =* 0.064; Tab. 2; Fig. 2).

**Figure 2.**
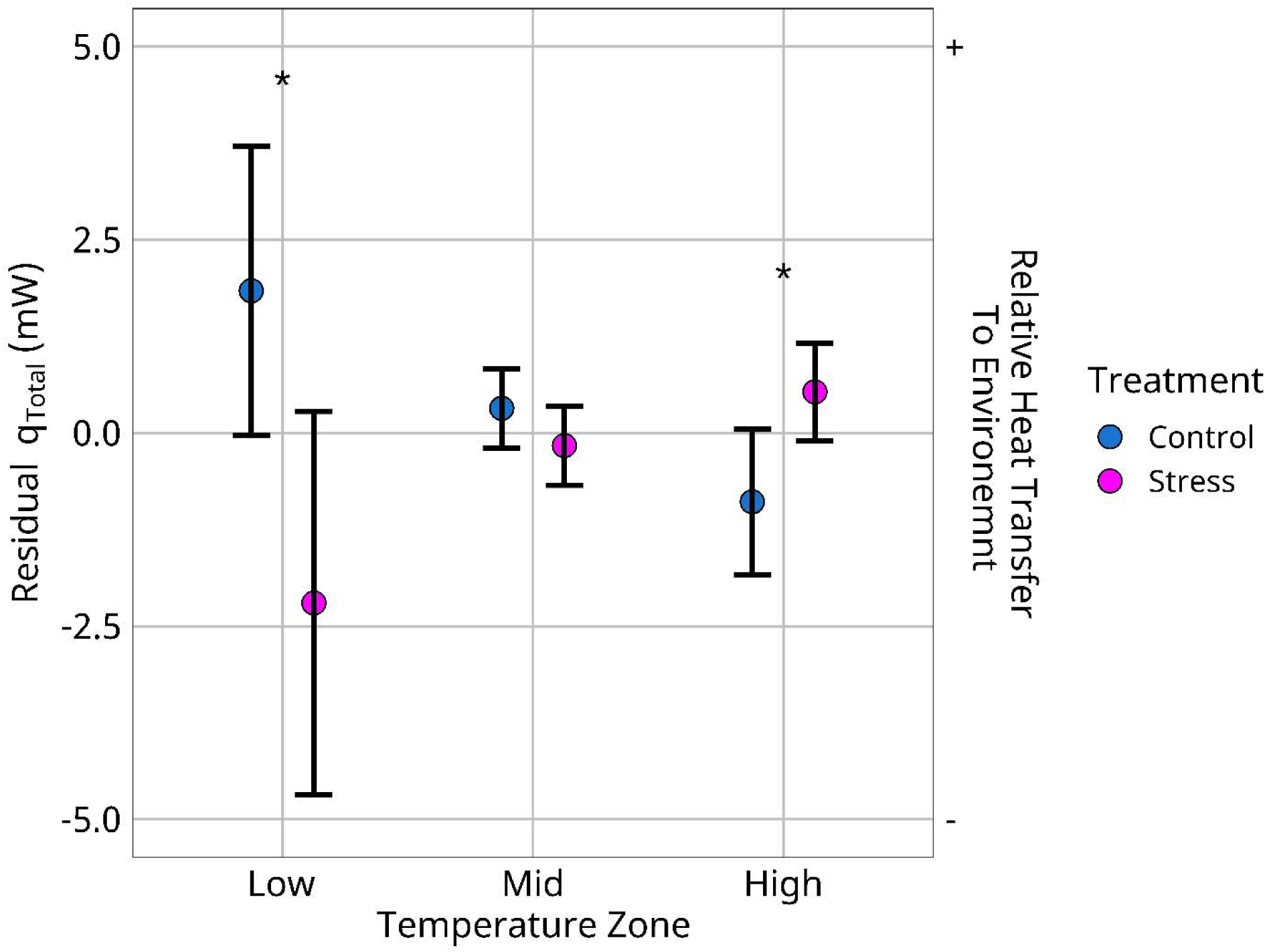
Residual heat exchange (q_Tot_) from the eye region of Black-capped Chickadees (n = 20) in response to repeated stressors, and across temperature zones. Residual q_Tot_ was calculated from a generalized additive model regressing q_Tot_ (mean per individual, per hour) against ambient temperature (mean *Ta* per hour; °C), then plotted again temperature zone. Low, mid, and high temperature zones represent *Ta* values below, at, and above thermoneutrality respectively (n_low_ = 111, n_mid_ = 710, n_high_ = 185; n_Total_ = 1006). Dots represent raw group means, and whiskers represent 95% confidence intervals around means. Blue colour represents control treatments, and pink colour represents stress treatments. Asterisks represent significant differences at an *a* of 0.05.

**Table 2.**
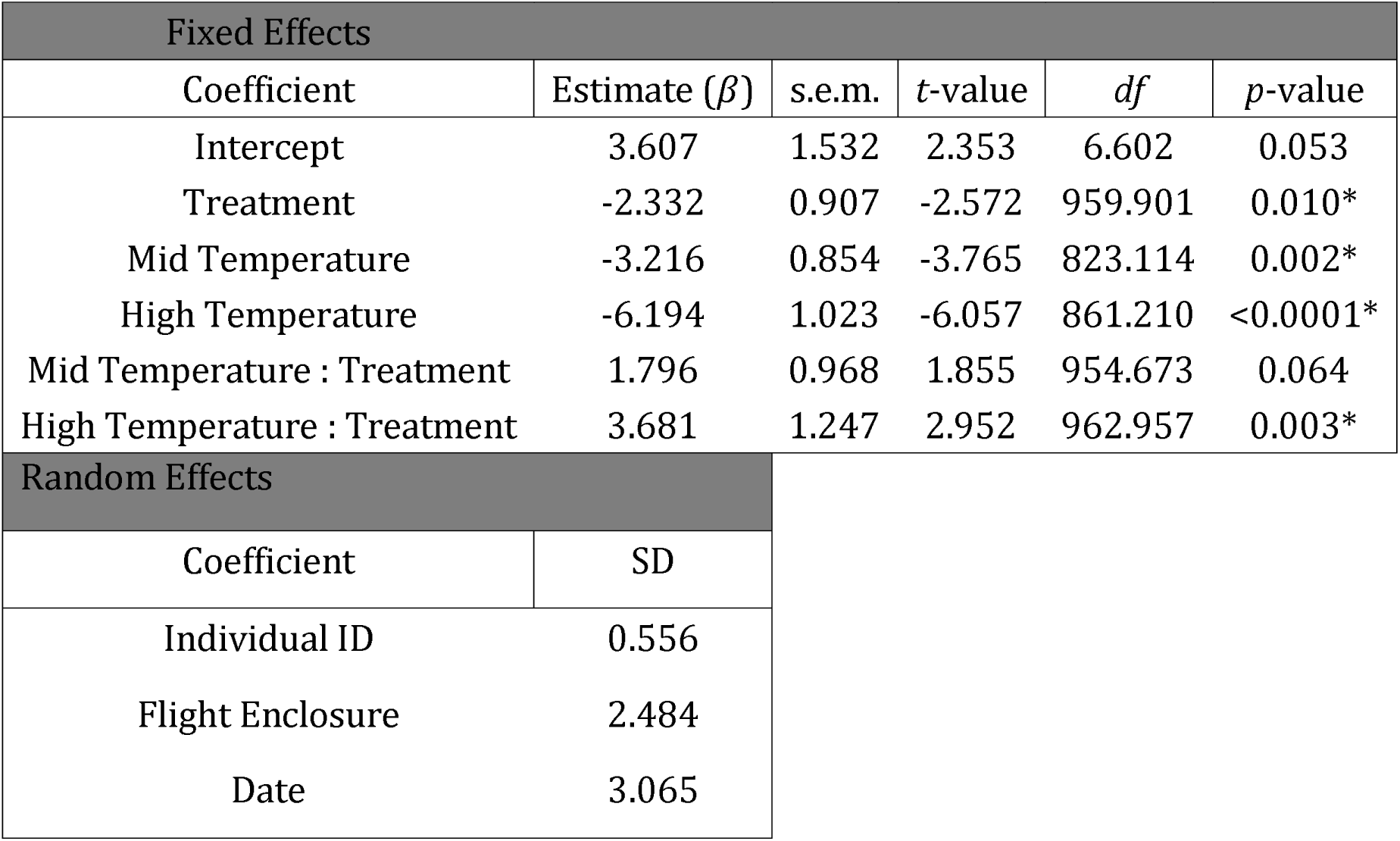
Results of an LMM testing the influence of temperature zones (below, at, and above thermoneutrality; “low”, “mid” and “high” respectively) and stress-exposure on total heat-loss from the eye regions of Chickadees. The low temperature and control categories are used as reference points for comparison of other categories and treatments. Standard errors (s.e.m.) are in reference to coefficient estimates (*β*). Asterisk (*) indicates significance ata = 0.05;n = 1006; n_Birds_ = 20; n_Days_ = 60.

| Fixed Effects |  |  |  |  |  |  |
| --- | --- | --- | --- | --- | --- | --- |
| Coefficient | Estimate ($\beta$) | s.e.m. | *t*-value | *df* | *p*-value | |
| Intercept | 3.607 | 1.532 | 2.353 | 6.602 | 0.053 |  |
| Treatment | -2.332 | 0.907 | -2.572 | 959.901 | 0.010* |  |
| Mid Temperature | -3.216 | 0.854 | -3.765 | 823.114 | 0.002* |  |
| High Temperature | -6.194 | 1.023 | -6.057 | 861.210 | <0.0001* |  |
| Mid Temperature : Treatment | 1.796 | 0.968 | 1.855 | 954.673 | 0.064 |  |
| High Temperature : Treatment | 3.681 | 1.247 | 2.952 | 962.957 | 0.003* |  |
| Random Effects |  |  |  |  |  |  |
| Coefficient | SD |  |  |  |  |  |
| Individual ID | 0.556 |  |  |  |  |  |
| Flight Enclosure | 2.484 |  |  |  |  |  |
| Date | 3.065 |  |  |  |  |  |

### Stress-induced changes in surface temperature, but not heat transfer, differ between sexes

Females and males significantly differed in their responses to stress exposure across *Ta*, as predicted by the Thermoprotective Hypothesis but not the Haemoprotective hypothesis. Specifically, eye region temperature in males displayed a more robust response to stress exposure than eye region temperature in females, according to *Ta* (n_females_ = 10, n_males_ = 10; *β*_males_ = 0.029, s.e.m. = 0.014, *t=* 2.138, *df=* 18, *p =* 0.046), with the slope between *Ta* and eye region temperature being steeper following stress exposure in males than females (Fig. 3). Interestingly, q_Tot_ from the eye region did not reflect patterns that were shown for eye region temperature, however. A global three-way interaction between sex, treatment, and temperature zone was not significant (F = 1.928, *df* = 799, *p =* 0.146) demonstrating that we could not detect sex-specific changes in heat transfer across temperature zones and treatment, probably owing to a small sample size (n_females_ =10, n_males_ =10).

**Figure 3.**
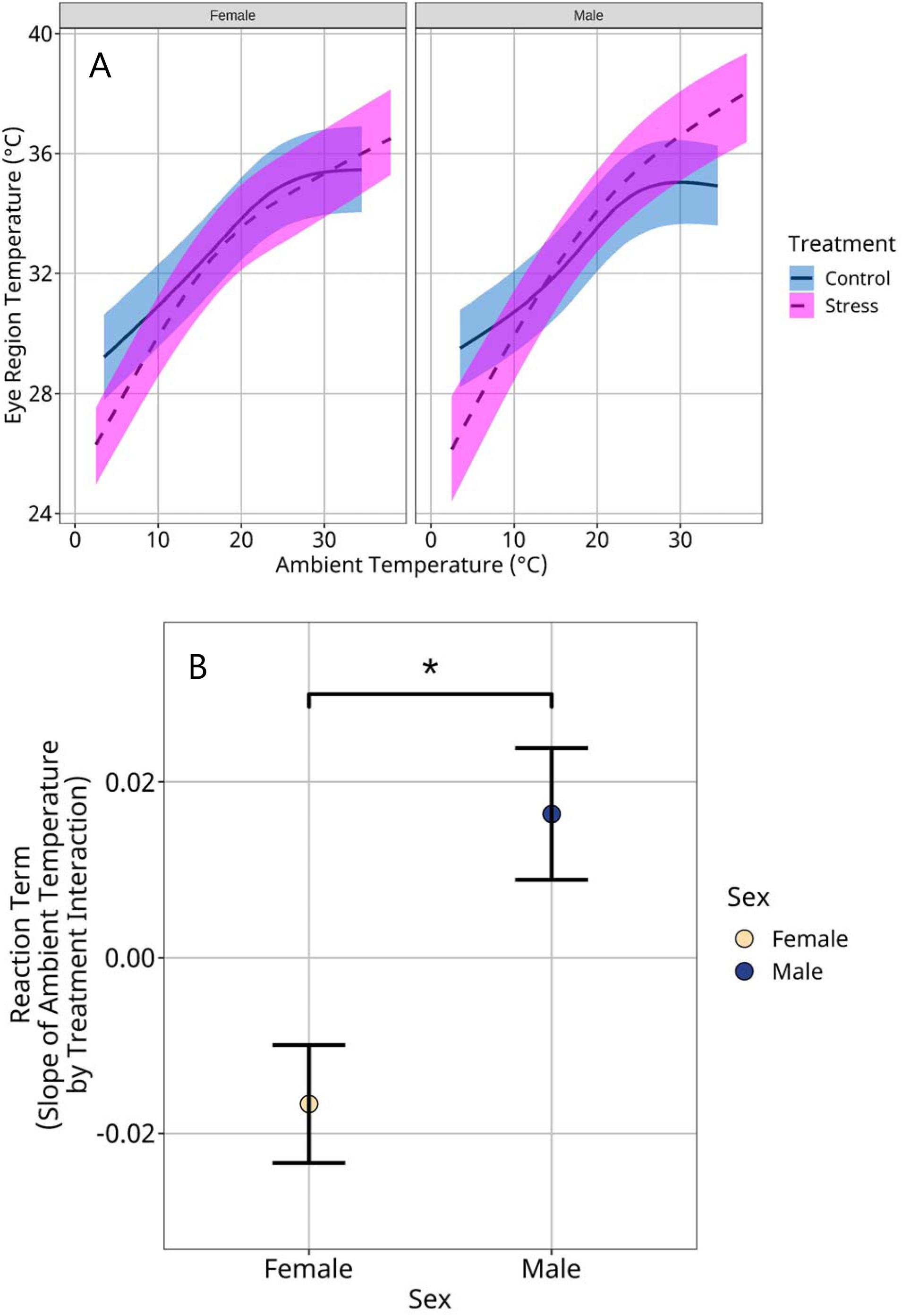
Sex-specific effects of repeated stress exposure on eye region temperature (°C) of Black-capped Chickadees (n_females_ = 10, n_males_ = 10) across a *Ta* gradient (°C). Eye temperature measurements were obtained by infra-red thermography across sixty days (n = 6431 images). **A |** Effect of sex on eye temperature according to *Ta* and treatment. Curves were calculated according to average treatment by *Ta* slope per *sex*, from GAMM output. Solid lines represent control groups, and dashed lines represent treatment groups. Blue and magenta shadows represent 95% confidence intervals around mean estimates for control groups and treatment groups respectively. **B |** Mean interactive effect of treatment and *Ta* on eye temperature, as compared between sexes. Mean interaction terms were derived from random slopes per sex, of a GAMM. Dots represent reaction term means, and whiskers represent 95% confidence intervals around means. Asterisks represent significant differences at an *a* of 0.05.

## Discussion

### Changes in surface temperature following stress exposure provide thermoregulatory advantages

Our results provide strong evidence for the Thermoprotective Hypothesis over the Haemoprotective Hypothesis, and lend support to direct functional significance of stress-induced changes in surface temperate. Indeed, as predicted by the Thermoprotective Hypothesis, we report a significant interaction between *Ta* and treatment on eye region temperature in our experimental population, where eye region temperature in stressed individuals was lower at low *Ta*, and higher at high *Ta* than that of control individuals (Fig. 1). Furthermore, individuals exposed to stressors conserved more heat at temperatures below thermoneutrality and dissipated more heat at temperatures above thermoneutrality with respect to controls (Fig. 2). These results demonstrate that changes in surface temperature experienced during stress exposure provide a thermoregulatory function across ecologically relevant temperature gradients, and likely contribute to lessening thermal burdens experienced during activation of a stress response. In our study, the effect of treatment on eye region temperature across *Ta* could not be explained by feeding behaviour or individual condition alone (SFigs 2-4), supporting direct physiological modification of surface temperature in response to stress exposure.

Although our reported patterns of q_Tot_ under stress are suggestive of a thermoregulatory adaptation, we recognize that our observed differences in q_Tot_ between stress-exposed and control individuals are small (mean Δq_Tot_ = 2.732 mW, or 0.937 J/g/hour; Fig. 2). Indeed, for a stress-exposed individual of average mass within our study population (10.5 grams), energetic savings accrued by such changes in q_Tot_ sum to approximately 1% of daily energetic expenditure, as estimated according to Karasov et al (1992). Given we assessed q_Tot_ across a small area of surface tissue alone (the eye region), however, our estimates of energy savings likely underestimate true energy savings associated with whole-body changes in surface temperature following stress exposure. In European Starlings (*Sturnus vulgaris*), for example, most dry heat transfer occurs at the ventral brachial area and legs (Ward et al, 1999), while in the Toco Toucan (*Ramphastos toco*), dry heat transfer is predominantly sanctioned to the bill (Tattersall et al, 2009). Combined heat transfer across the legs, bill, and eye region in our birds almost certainly exceed that across the eye region alone, if thermal responses to stress exposure at the legs and bill mimic those observed at the eye region (but see Bech and Midtgård, 1981). Indeed, Herborn et al (2015) reported a concomitant change in wattle, comb, and eye temperature following stress exposure in the Domestic Chicken, demonstrating that surface tissues are unlikely to respond in isolation to stress exposure. In this study, however, we were unable to estimate q_Tot_ across the legs and bill of individuals because each structure could not be readily observed within our thermographic images.

Beyond providing direct thermoregulatory advantages, stress-induced changes in surface temperature may also yield indirect advantages with respect to water retention. In small birds, active heat dissipation is predominantly achieved by evaporative cooling at cutaneous tissues in the respiratory tract (via panting; Tieleman and Williams, 2002). Cooling by this mechanism, however, can provide significant water-loss that may be challenging to sustain. Black-capped Chickadees, for example, have been shown to lose up to 400 mg of water per hour by evaporative cooling at *Tas* above thermoneutrality (4% of their body mass; Rising and Hudson, 1974). Estimates for other temperate species are comparable, with Red Breasted Nuthatches (*Sitta canadensis*) losing up to 260 mg of water per hour by evaporative cooling at high *Ta* (2.6% of their body mass; Mugaas and Templeton, 1970). As evaporative heat-loss at surface tissues is thought to be negligible in contrast to respiratory tissues (Bernstein, 1971b), heat dissipation at surface tissues by radiation or conduction therefore provides significant water-retentive advantages when compared with cooling by panting. In desert-dwelling birds, for example, total evaporative water loss may be reduced by 50% when employing methods of dry heat exchange (i.e. peripheral hyperthermia) rather than evaporative cooling (Tieleman and Williams, 1999). Under stressful conditions, water stores may already be compromised (i.e. by stress-induced defecation; Jones et al, 1995; Haas et al, 2010) and the ability to seek and acquire water may be over-ruled by combative or avoidance behaviours. For this reason, elevations in heat transfer from surface tissues of stress-exposed Chickadees could also reflect the outcome of a thermoregulatory trade-off at high *Ta*, where dissipation of heat generated during a stress response, albeit small, is balanced by a pressure to retain water stores.

### Male Surface Temperature is More Responsive to Stress Exposure than Female Surface Temperature

In our study, the effect of treatment on the relationship between eye region temperature and *Ta* was significantly greater in males than in females (Fig. 3). This trend was consistent with predictions of the Thermoprotective Hypothesis, as male Black-capped are thought to have both lower thermal conductance and heat dissipation capacity than females (Aschoff, 1981; Porter and Kearney, 2009), owing to their larger body size (Foote et al, 2010). To our knowledge, no other studies have tested the effect of sex differences in surface temperature responses to stress exposure. Interestingly, our analyses did not detect differences in the effect of stress exposure on q_Tot_ between sexes, however, a low sample size (n_females_ = 10, n_males_ = 10) probably contributed to a lack of statistical clarity (*p* = 0.146).

Given our experiment was initiated at the onset of the breeding season for Black-capped Chickadees (April; Foote et al, 2010), it is possible that our observed differences in surface temperature under stress exposure are a consequence of differential thermal windows between sexes. Female Black-capped Chickadees, unlike males, display incubation behaviour, and brood-patch development is consequentially limited to them alone (Foote et al, 2010). Female brood patch development begins in April at a comparable latitude to our field location (Odum, 1941; Cooper and Voss, 2013); approximately the same time-point at which our experiments were initiated. Although we did not observe abdominal defeathering in our birds during experimentation, females may have displayed greater abdominal vascularization inherent to brood patch development than males (Hinde, 1961; Etche et al, 1979; but see Bailey, 1952), allowing them enhanced heat dissipation capacity at high temperatures (Hill et al, 2014; Nilsson and Nord, 2018), and leveling the slope between *Ta* and eye region temperature in stress treatments, above thermoneutrality.

Notably, a more robust change in surface temperature following stress exposure in males could reflect a stronger selective pressure for thermoregulation in this sex, particularly when both *Ta* and metabolic heat production are high (i.e. during activation of a stress response in warm environments). Differences in thermal fertility limits between sexes have recently been argued as an emerging trend across species, with males typically displaying higher thermal sensitivity than females (Iossa, 2019). Such sex differences may be exacerbated in taxa where males have internal testes (i.e. birds). Sex-specific thermal limits to fertility could exert a stronger selection pressure for thermoregulatory capacity, or heat dissipation capacity, in male Chickadees at high *Ta* when compared with female Chickadees. Higher eye region temperature in males Chickadee exposed to both high *Ta* and perceived stressors may therefore be explained by adaptations to dissipate more heat (that is generated by heat exchange with the environment, and as a byproduct of mounting a physiological and behavioural stress response) at the level of surface tissues when compared to females.

Beyond differences in thermal windows and heat dissipation capacity, the disparity in stress-induced changes in surface temperature between sexes may also be explained by differences in autonomic and steroidal pathways underpinning the thermal response to stress. In birds, males typically display a more robust activation of the hypothalamic-pituitary-adrenal axis following stress exposure (Silverin, 1998; Marin et al, 2002; Madison et al, 2008; Wada et al, 2008). Across vertebrates, HPA axis activation is thought to both enhance sympathetic function (Fisher et al, 1982), and sensitize the body to catecholamine availability (Fisher et al, 1982; Krakoff, 1988), thereby exacerbating the tachycardic, tachypneic, and haemodynamic response to a perceived stressors (reviewed in Sapolsky et al, 2000). In our study, differences in thermal response curves between sexes (across *Ta* and treatments) may simply be explained by exacerbated vasoconstriction at the periphery in males below thermoneutrality (Fig. 3), and elevated total heat production in males above thermoneutrality (Fig. 3), each as a consequence of enhanced physiological stress-responsiveness (i.e. tachycardia, tachypnea, and avoidance behaviour; Cabanac and Aizawa, 2000; Cabanac and Guillemette, 2001; Greenacre and Lusby, 2004; Long et al, 1990a).

## Summary

Here, we proposed a new hypothesis stating that changes in surface temperature following exposure to stressors hold direct functional significance by offsetting thermoregulatory costs experienced during activation of a stress response (the Thermoprotective Hypothesis). This hypothesis contrasts dominant theory which argues that changes in surface temperature following stress exposure are a mere consequence of haemetic redistribution and hold no direct functional significance in themselves. Our results strongly support the Thermoprotective Hypothesis, and consequently provide evidence for functional significance of stress-induced changes in surface temperature. Indeed, we show that in a temperate endotherm, stress-induced changes in surface temperature influence dry heat transfer to the environment and allow individuals to conserve more heat at low *Ta*, and dissipate more heat at high *Ta* when experiencing prolonged stress exposure. These findings emphasize a seldom-recognized influence of the thermal environment in shaping physiological responses to stress in vertebrates, but in doing so, raise questions about the suitability of such responses within the context of a changing global climate.

## Supporting information

Supplemental Methods and Results

## Acknowledgements

We thank Lianne Ralph for exceptional patience and field assistance, Glenn Tattersall for thermographic advice, Simon Tapper and Robby Marrotte for statistical input, Jan-Åke Nilsson and Andreas Nord for theoretical conversations, James Quinn for his generous contribution of laboratory space, and Kimberley Tasker and Tyler Maksymiw for their assistance in aviary construction. Additionally, we thank the staff and avian banding team at the Ruthven Park National Historic Site for their immense support in experimental procedures.

