## Supplemental Methods and Results for "Evidence that stress-induced changes in surface temperature serve a thermoregulatory function"

### Supplemental Material

#### Methods

***Statistical Analysis***

Exposure to repeated stressors significantly influenced feeding behaviour in our experiment (see "Results"; Tab. 2). We therefore tested whether this effect of treatment on feeding behaviour (as quantified by feeding score; see "Methods") was sufficient to explain our observed differences in eye temperature across ambient temperature (*Ta*) between treatment groups (see "Results"; Tab. 1). We therefore asked whether eye temperature was correlated with feeding score, when controlling for the influence of ambient temperature on both parameters. To address this question, we first averaged eye temperature for each individual per hour, per day, then calculated average *Ta* during these respective time period. We then quantified the relative difference between observed eye temperature means and those expected at any given *Ta* (regardless of treatment) by constructing a simple generalized additive model ("GAM"; R package mgcv ) with mean eye temperature as the response variable, and mean *Ta* as a smoothed predictor (cubic regression spline with four knots), then extracting residual eye temperature from this GAM. Our previous results (see "Results"; Tab. 1) show that the direction of the effect of treatment on eye temperature varies according to *Ta*. We therefore calculated the absolute values of the eye temperature residuals to create a directionless metric of distance between observed eye temperature means, and expected eye temperature means for a given *Ta*.

To test whether differences in feeding rate, but not treatment, could explain discrepancies between mean eye temperature for a given individual and that expected for a respective *Ta*, we constructed a generalized linear mixed effects model ("GLMM"; R package cplm ) with residual eye temperature (absolute values, as described above) as the response variable, and feeding score and treatment (binomial factor) as fixed effect predictors. Time (UT1 Hour) was also included is a fixed effect to account for circadian rhythms in body temperature, as well as an interaction between time and treatment to account for significant interaction between each variable observed in our previous model (see "Results"; Tab. 1). Residual effects of individual identity, flight enclosure identity, and date were initially accounted for by including each parameter as a random intercept, however, individual identity explained zero variance in our model and was therefore removed. A Tweedie distribution was assumed for our model as our response variable was right-skewed, non-negative, and a non-integer. Our final model included 581 observations across 50 days, with all four flight enclosures considered. Alpha values were set to 0.05 for all analyses.

#### Results

Feeding score was not significantly correlated with residual eye temperature ($\beta$ = -0.001, *t* = -0.491, *df* = 581, *p* = 0.312; SFig. 1), but was significantly influenced by treatment, with stressed individuals displaying high residual eye temperature value ($\beta$ = 1.444, *t* = 2.811, *df* = 581, *p* = 0.003; SFig. 2). Similar to our previous results ("Results", Tab. 1), time alone was not significantly correlated with eye temperature ($\beta$ = 0.041, *t* = 1.105, *df* = 581, *p* = 0.135), however, a significant interaction between time and treatment was observed ($\beta$ = -0.137, *t* = -2.794, *df* = 581, *p* = 0.003).

#### Figure Legends

**Supplemental Figure 1 | Residual eye temperature (**$\mathbf{}^{\boldsymbol{\circ}}$**C) of Black-capped Chickadees (n = 20) according to feeding rate, per individual.** Eye temperature residuals (absolute values, raw) were extracted from a generalized additive model with mean eye temperature per individual, per hour and day, regressed against respective mean ambient temperatures (Ta, ${}^{\circ}$C), per hour and day. Feeding Score represents the combined effects of feeding visits (visits/hour) and time spent feeding (seconds/hour), as derived from a principle component analysis and normalized between 0 and 100. Observations (n = 581) were derived from 50 days, across four flight enclosures.

**Supplemental Figure 2 | Residual eye temperature (**$\mathbf{}^{\boldsymbol{\circ}}$**C) of Black-capped Chickadees (n = 20) according to experimental treatment**. Eye temperature residuals (absolute values, raw) were extracted from a generalized additive model with mean eye temperature (${}^{\circ}$C) per individual, per hour and day, regressed against respective mean ambient temperatures (Ta, ${}^{\circ}$C), per hour and day. Dots represent marginal means of residual eye temperature for each treatment, as derived from a mixed-effect model. Whiskers represent 95% confidence intervals around mean estimates. Observations (n = 581) were derived from 50 days, across four flight enclosures.
