## Supplementary figures and images for "Evidence that stress-induced changes in surface temperature serve a thermoregulatory function"

### Supplemental Figure 1

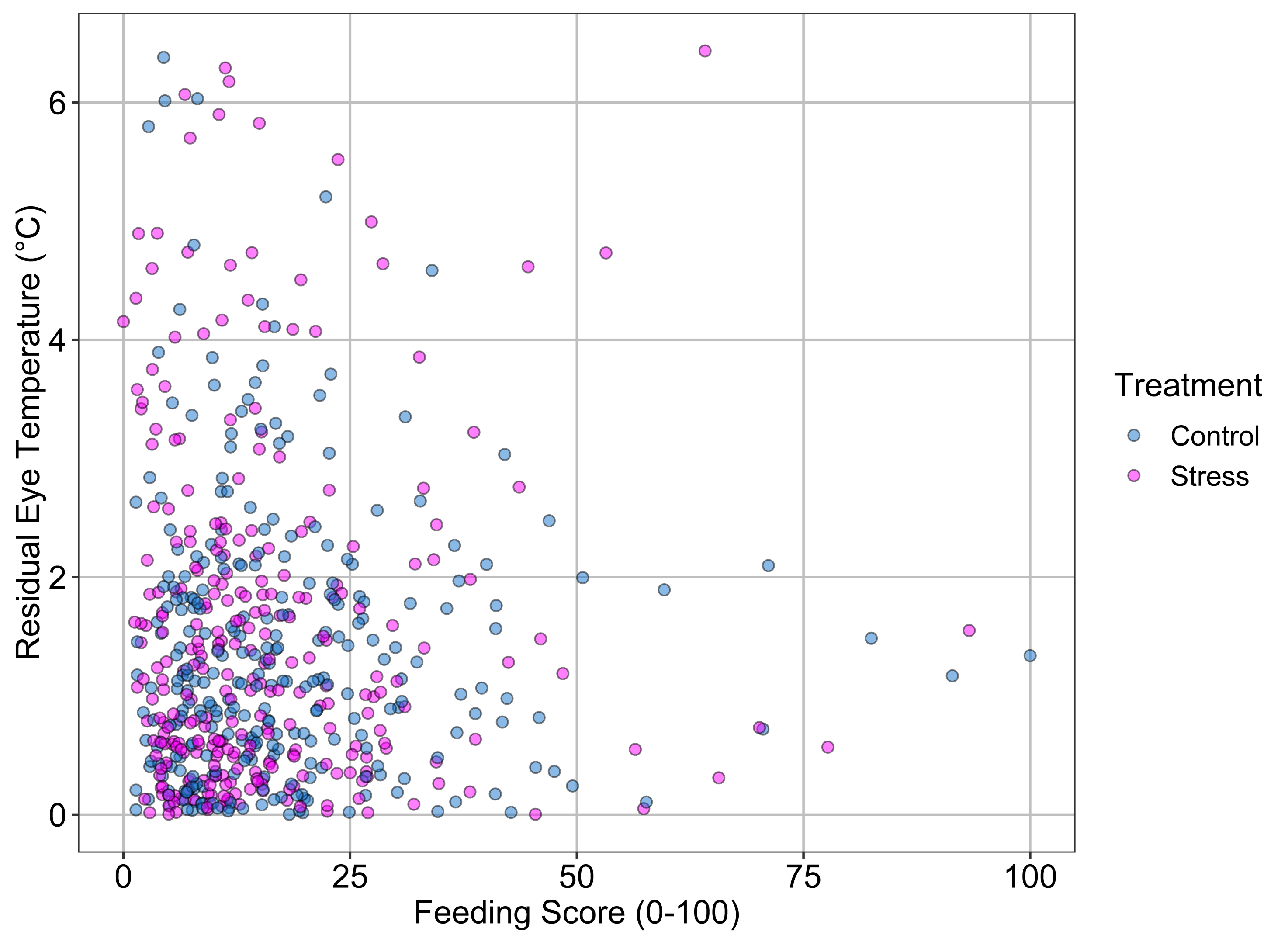

### Supplemental Figure 2

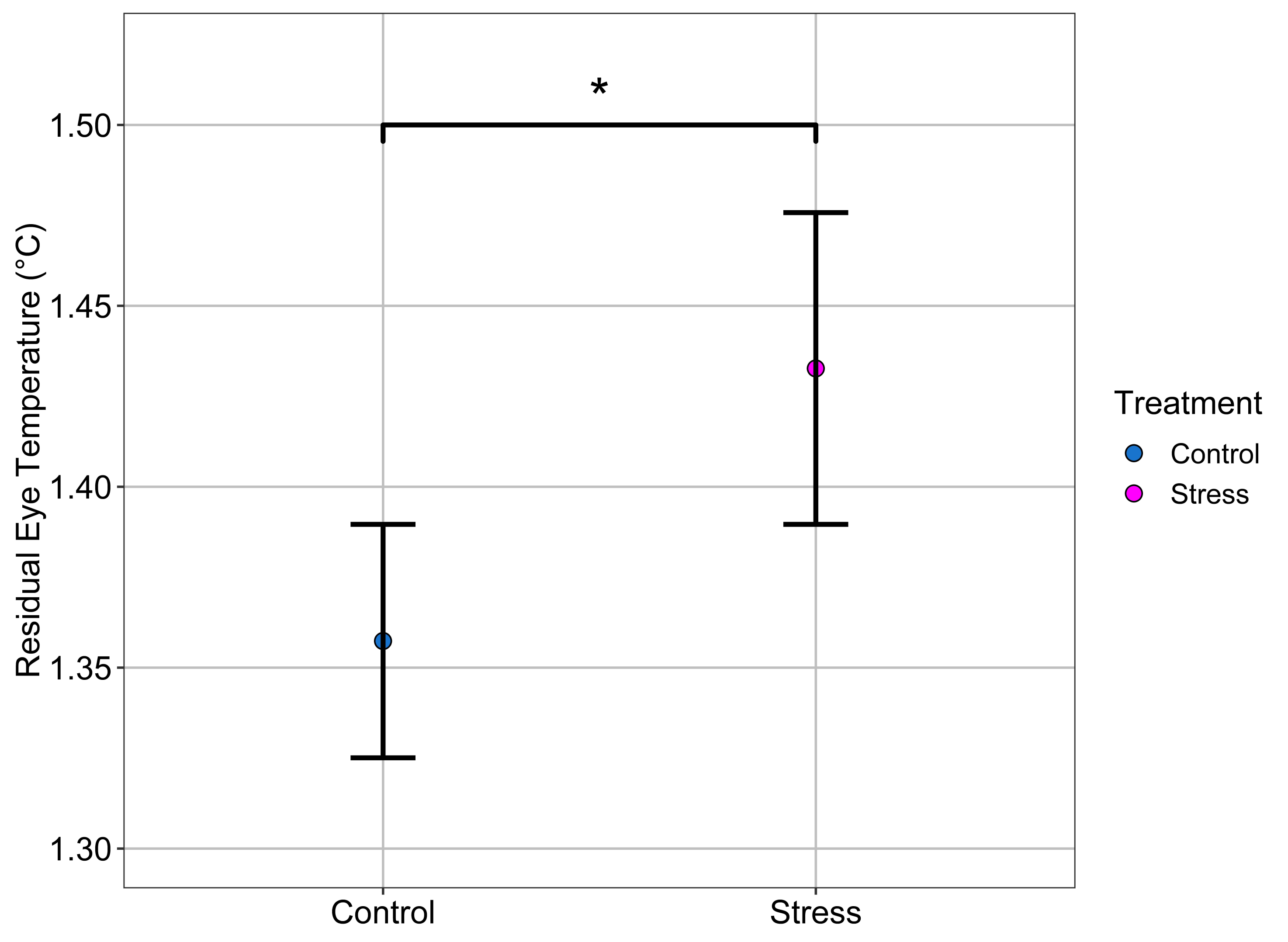
